# Isolation and Characterization of Extracellular Vesicles and Non-Vesicular Extracellular Particles from Mouse Tissues

**DOI:** 10.1101/2025.03.02.641099

**Authors:** Marta Garcia-Contreras, Worawan B. Limpitikul, Saumya Das

## Abstract

Extracellular vesicles (EVs) and non-vesicular extracellular particles (NVEPs) have been recently discovered as part of the diverse components of the secretome. EVs and NVEPs have been shown to differ not only in size and morphology but also in their cargo and biological functions. NVEPs are a newly discovered group of multimolecular assemblies with potential roles in physiological and pathological states. However, very little is known about this new class of particles. Here, for the first time, we present a method for simultaneously isolating and characterizing EVs and NVEPs from primary mouse tissues.

## Introduction

Communication between different cell types is necessary for maintaining tissue homeostasis. The secretomes (proteins, peptides, extracellular vesicles, non-vesicular extracellular particles, etc.) of different cell types have emerged as an important mediator of cell-to-cell signaling. As a part of the cell secretomes, extracellular vesicles (EVs) and non-vesicular extracellular particles (NVEPs), are secreted from various cell types and have been found in biological fluids^1^.

EVs are lipid bilayer membrane vesicles released from cells. EVs can be classified into two subtypes based on size: large (l-EVs) that are >200nm in diameter and small EVs (s-EVs) that are <200nm in diameter. EVs are enriched in tetraspanin markers (CD63, CD9, and CD81), and contain protein cargo including syntenin-1, tumor susceptibility gene 101 (Tsg101), Programmed cell death 6-interacting protein (Alix), or Flotillin-1^2^.

In contrast, NVEPs are a group of recently discovered multimolecular assemblies, that are not enclosed in a lipid bilayer and are released from most of cell types^3^. NVEPs have been classified into two subpopulations, exomeres and supermeres, based on their isolation by a differential centrifugation method^4,5^. Exomeres and supermeres are small NVEPs <50 nm in diameter. They differ in size, content (nucleic acids and proteins), and biological functions^1,6,7^. While both exomeres and supermeres are enriched in metabolic enzymes, such as AGO2 (Argonaute RISC Catalytic Component 2), lactate dehydrogenase A (LDHA), Transforming growth factor-beta-induced protein ig-h3 (TGFBi) and Heat shock 70 kDa protein 13 (HSPA13), supermeres are enriched in RNA-binding proteins^1^. Additionally, supermeres are enriched in disease-related cargo such as amyloid precursor protein (APP) and amyloid beta precursor like protein 2 (APLP2) in Alzheimer’s disease; and angiotensin converting enzyme (ACE) and ACE2 in cardiovascular disease^3^ and SARS-CoV-2^8^, making them a potential source of novel biomarkers. Nonetheless, very little is known about NVEPs.

In this manuscript, we present a method to isolate simultaneously EVs and NVEPs from primary mouse tissues for the first time. This protocol might be applied for investigating their role in pathogenesis or physiology of different tissues.

## Methods

### Animal Protocol and Maintenance

This study was conducted on 6-week-old C57BL/6J male mice (#000664 The Jackson Laboratory). All animal experiments were performed according to the approved Massachusetts General Hospital Administrative Panel on Laboratory Animal Care (APLAC). Mice were maintained in 12 h light/dark cycles and were fed with a standard irradiated rodent chow diet.

### Tissue preparation and processing

Mouse brain, quadriceps, and epidimal white adipose tissue (eWAT) were dissected and placed in 10-cm a petri dishes with phosphate buffered saline (PBS) (Catalog number 10010023, Gibco). Then, the tissues were cut into smaller pieces while submerged in tissue culture media (DMEM/F12 (Catalog number 11330032, Gibco) or RPMI 1640 medium (Catalog number 11875135, Gibco) supplemented with 1% penicillin-streptomycin (Catalog 15140163, Thermo Scientific) for brain and quadriceps tissues respectively, and RPMI 1640 medium (Catalog number 11875135, Gibco) supplemented with B27 supplement (Catalog number 17504044, Thermo Scientific) and 1% penicillin-streptomycin (Catalog number 15140163, Thermo Scientific) for eWAT in a well from a 6-well plate (Catalog number 353046, Corning) and incubated either for a minimum of 4 hours or overnight in an incubator (37°C, 5% CO_2_). Culture medium was then collected and passed through a 70-µm cell strainer (Catalog number 352350, Corning) to remove small pieces of tissue. We further removed smaller debris by placing the media in a 15-ml tube and centrifuge at 2,000xg for 20 minutes at 4 °C. Then the supernatant (conditioned media) was transferred to a new 15-ml tube and stored frozen at −80 °C until further processing.

### Extracellular Vesicles and Non-Extracellular Vesicle Particles Isolation

We adapted established methods for isolating EVs and NEVPs from cell culture and human biological fluids^9^. We centrifuged thawed conditioned media at 10,000xg for 40 minutes at 4 °C. The pellet was resuspended in 200µl of PBS containing 20 mM HEPES (PBS-H) to obtain large-EVs. The supernatant was further centrifuged at 167,000xg for 4 hours at 4 °C. The pellet after this centrifugation was resuspended in 200µl of PBS-H to obtain small-EVs while the supernatant was further centrifuged at 167,000xg for 16 hours at 4 °C. Afterwards, the resultant pellet was resuspended in 200µl of PBS-H to obtain exomeres while the supernatant was further centrifuged at 367,000xg for 16 hours at 4 °C. The final pellet was resuspended in 200µl of PBS-H to obtain supermeres.

### Protein quantification and Western blotting

Protein was quantified by a BCA Protein Assay (Catalog number 23227, Thermo Scientific) following manufacture’s protocol. For western blots, proteins were lysed in RIPA buffer (Catalog number 89900, Thermo Scientific) supplemented with Protease and Phosphatase Inhibitor Cocktail (Catalog number 78441, Thermo Scientific). Protein lysates were resuspended in 4× Laemmli protein sample buffer (Catalog number 1610747, Bio-Rad) and denatured at 95 °C for 5 min. The denatured protein samples were then run on Tris/Glycine/SDS pre-cast stain free polyacrylamide gels (Catalog number 5678094, Biorad) for 70 minutes at 110 V. Proteins were then transferred onto polyvinylidene fluoride membranes (Catalog number 1704273, Biorad) using a semidry transfer method following manufacture’s protocol. The membranes were blocked in 5% BSA in Tris-buffered saline containing 0.1% Tween 20 (TBS-T) for 1 h. Next, the membranes were incubated with primary antibodies (Table 1) in blocking buffer overnight at 4 °C, washed 3 × 10 min in 0.1% TBST, and incubated with the appropriate horse radish peroxidase-conjugated secondary antibody (Table 1) in blocking buffer for 1 h at room temperature. The membranes were then washed in 0.1% TBS-T (3 × 10 min) and developed with Supersignal West Femto Maximum Sensitivity Chemiluminescent Substrate (Catalog number 34095, Thermo Fisher) in a 1500 iBright Imaging system (Thermo Fisher Scientific)

**Table 1.** Antibody List

| Antibody | Source | Catalog Number | Dilution |
| --- | --- | --- | --- |
| Anti-CD81 | Thermo Fisher Scientific | MA5-32333 | 1:1000 |
| Anti-CD63 | Abcam | ab216130 | 1:1000 |
| Anti-LDHA | Cell signaling | 2012S | 1:1000 |
| Anti-beta actin | Santa Cruz | A5316 | 1:500 |
| Anti-TMEM119 | Genetex | GTX639083 | 1:1000 |
| Anti-Adiponectin | Cell Signaling | 2789S | 1:1000 |
| Anti-TGFBi/BIGH3 | Proteintech | 10188-1-AP | 1:1000 |
| Anti-AGO2 | Abcam |  | 1:500 |

### Microfluidic resistive pulse sensing (MRPS)

We performed microfluidic resistive pulse sensing (MRPS) (nCS1, Spectradyne LLC, USA) on small EVs. EVs were re-suspended either at 1:50 or 1:100 dilution, and 10μl were loaded into the corresponding cartridge of 0.1% Tween-PBS filtered through a 0.02 μm filter (Catalog 6809-3002, Cytiva). All samples were measure using C-2000 cartridges: ∼ 250–2000 nm or C-400 cartridges ∼ 65–400 nm particle size range. MRPS Spectradyne measurements are plotted to show particle size (concentration/ml) on Y-axis and particle diameter (nm) on the X-axis.

### Statistical analyses

Statistical analyses were performed using GraphPad (Prism, Version 10.0.2). A two-sided Student’s t-test (adjusted for unequal variance where applicable) was used to compare the difference in means across sample groups. P values of < 0.05 were considered statistically significant in all analyses.

### Software

All illustrations were created with BioRender.com.

## Results

### Size distribution of large and small EVs isolated from primary tissues

EVs and NVEPs were isolated from the brain, quadriceps, and epididymal white adipose tissue (eWAT) using differential centrifugation, which allowed for the separation of different vesicle populations based on their size and density. Two distinct time points were tested during the isolation process (4 hours or overnight incubation) to assess the yield, to evaluate how the efficiency of particle recovery changes over time. To further characterize the isolated particles, the size distribution of small extracellular vesicles (s-EVs) was analyzed using microfluidic resistive pulse sensing (MRPS).Small EVs (s-EVs) showed the expected size range: quadriceps-derived s-EVs ∼ 65-280 nm, brain-derived s-EVs ∼ 65-300 nm, and eWAT-derived sEVs ∼ 65-274 nm. Particle concentration of quadriceps-derived s-EVs is 1.0±0.3 x10^11^ (4-hour incubation) and 1.2±0.3 x10^11^ (overnight incubation) particles/ml, brain-derived s-EVs 2.3±0.5 x10^11^ (4-hour incubation) and 3.9 ±0.5 x10^11^ (overnight incubation) particles/ml, and eWAT-derived s-EVs 2.6±03 x10^10^ (4-hour incubation) and 2.7 ±0.1 x10^10^ (overnight incubation) particles/ml.

However, it is important to note that the size distribution of exomeres and supermeres, which are smaller and larger subtypes of EVs, respectively, could not be analyzed using the MRPS method. This is due to the limitations of MRPS in detecting particles outside the size range that can be accurately measured by this technique. Consequently, additional methods may be required to fully characterize the size distribution of these particle subtypes. Further investigations are needed to explore alternative approaches that can provide insights into the characteristics of exomeres and supermeres.

### Protein content of EVs and NVEPs

We next analyzed the protein concentration of EVs and NVEPs isolated from various tissues using a BCA Protein Assay. In the case of the quadriceps (Quad) and epididymal white adipose tissue (eWAT) primary tissues, we observed that overnight incubation resulted in a notable increase in the protein content for small extracellular vesicles (s-EVs) and exomeres. This suggests that the extended incubation period may enhance the yield of these particle types. Interestingly, despite the increase in protein content for s-EVs and exomeres, overnight incubation did not increase the protein amount for large extracellular vesicles (l-EVs) and supermeres in eWAT or Brain. In contrast, for brain tissue, overnight incubation did not lead to a significant increase in the protein content for any of the EV or NVEP groups.

### EV and NVEP marker characterization

To characterize the isolated particles, we characterized canonical EV markers (CD81 and CD63) and NVEP markers (AGO2 or LDHA), as well as tissue specific/enriched proteins (ADIPOQ for eWAT, TMEM119 for brain and ß-actin for quadriceps). CD81 and CD63 were found in EVs and exomeres. AGO2 was found to be enriched in EVs and exomeres, and LDHA in exomeres and supermeres for all tissues. Interestingly, tissue specific proteins were enriched in s-EVs and exomeres for all tissues.

## Conclusion

In this study, we successfully developed a method for isolating and characterizing extracellular vesicles (EVs) and non-vesicular extracellular particles (NVEPs) from primary mouse tissues - brain, quadriceps, and epididymal white adipose tissue (eWAT). The method effectively separates two classes of EVs, large and small, along with two distinct subpopulations of NVEPs, exomeres and supermeres. Using differential centrifugation, we were able to obtain purified particle fractions from each tissue adapting a previous protocol^4^. Previous protocols described the used of differential centrifugation to isolate EVs and NVEPs from cell culture media, human plasma and only exomeres from tissues^10,11^. Our adapted protocol, however, demonstrates for the first time the isolation EVs and NVEPs, including supermeres which has never being isolated before from tissues, using differential ultracentrifugation. Additionally, this Is the first study that examines different time points to determine the optimal conditions for mouse tissue isolation of EVs and NVEPs. Small EVs (s-EVs) were further analyzed for size distribution using microfluidic resistive pulse sensing (MRPS). The size distribution data confirmed that s-EVs were within the expected size ranges, consistent with prior literature, while the exomeres and supermeres were not measurable with MRPS due to their smaller size. This highlights a limitation in current methodologies for fully characterizing NVEPs, in the smaller size range. The particle concentration was similar for 4 hours and overnight incubations for all tissues. However, we observed tissue-specific differences in protein content after incubation, particularly in the quadriceps and eWAT tissues, where overnight incubation increased protein yields for s-EVs and exomeres. For brain-derived particles there was lack of a notable change in protein concentration, these findings suggests that incubation time or tissue-specific factors may play a role in the recovery of certain particle types. Further investigation is needed to understand the biological and methodological factors underlying these differences.

We further characterized EVs and NVEPs by immunoblotting. We found typical markers in the EV and NVEP fractions for all tissues, and for eWAT and Quad we saw an increased in expression after overnight incubation. AGO2 that was found in EVs and exomeres, it is still not clear to be an EV marker or a contaminant from the isolating methods^12^. Although tetraspanins CD81 and CD63 are known to be present in EVs there is a tissue distinct distribution as previously shown^13^, we found almost no expression of CD81 in exomeres in brain. But for all tissues there is an increased on CD81 expression after overnight incubation. Overall, our work presents a comprehensive and reproducible approach for isolating EVs and NVEPs from various tissues, providing a valuable tool for future studies exploring the role of these particles in physiology and disease. The differential isolation of EVs and NVEPs also opens new avenues for studying their respective cargos, including proteins, nucleic acids, and disease-related biomarkers, in the context of tissue-specific functions. Moving forward, refining the isolation methods and incorporating additional characterization techniques will be crucial for uncovering the full spectrum of biological roles these particles play in health and disease.

## Acknowledgments

The authors would like to thank the members of Das Lab

## Author contributions

M.G.C and W.L conceived the study; designed the experimental methodology; performed experiments; analyzed, interpreted, and visualized the data and wrote the manuscript. S.D supervised the research and edited the manuscript.

## Competing interests

SD is a founding member and has equity in Thryv Therapeutics and Switch Therapeutics. He also consults for Thryv Therapeutics. The rest of authors declare no competing interests.

## Figure Legends

**Figure 1:**
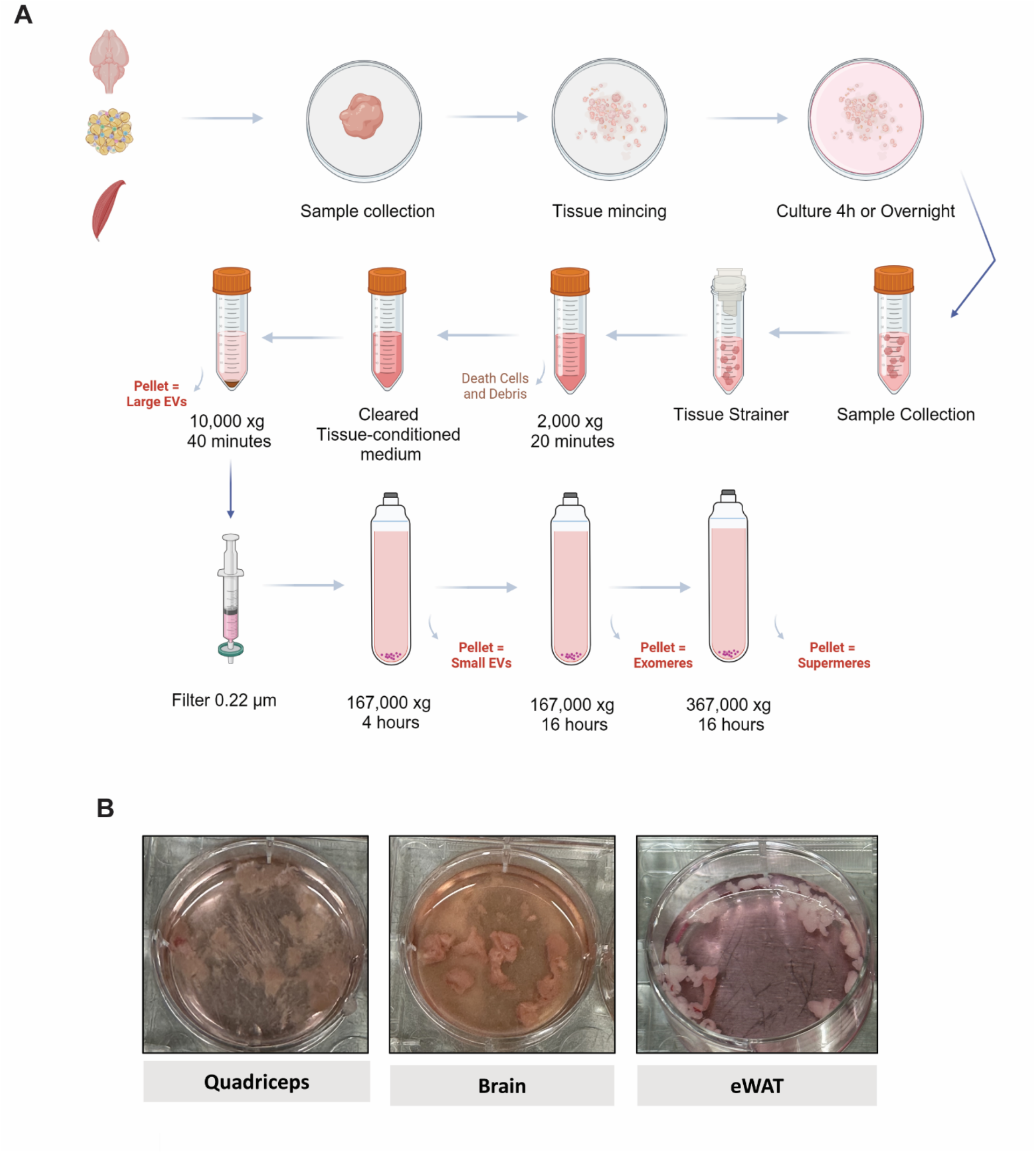
Overview of the Isolation protocol (**A**) Workflow of the isolation protocol for large EVs, small EVs, exomeres and supermeres (**B**) Representative images of the dissected tissues in culture.

**Figure 2:**
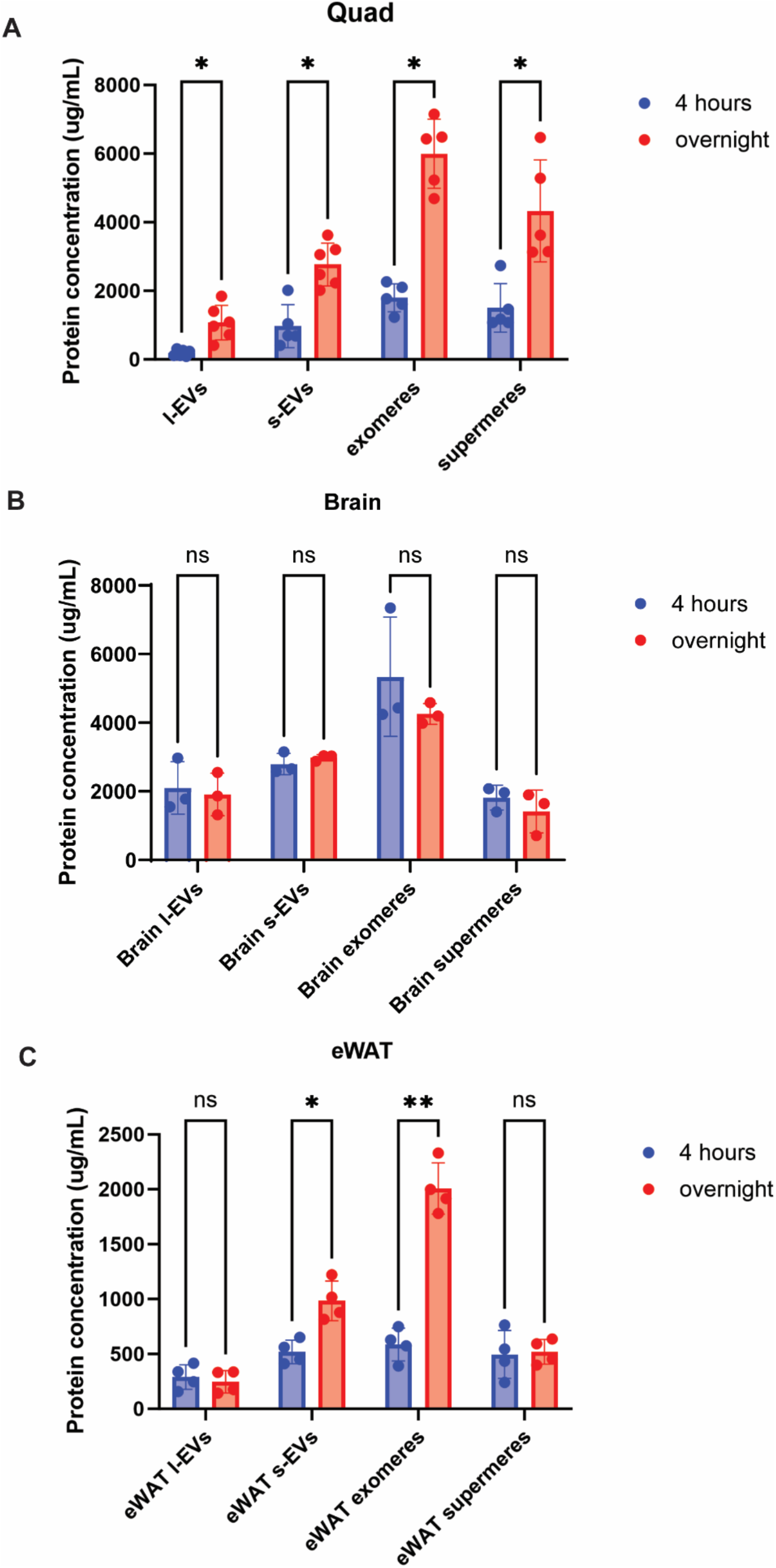
Comparison of protein concentration for EV and NVEPs-derived from Brain, Quad and eWAT tissue. Data is represented as (Mean ± SD).

**Figure 3:**
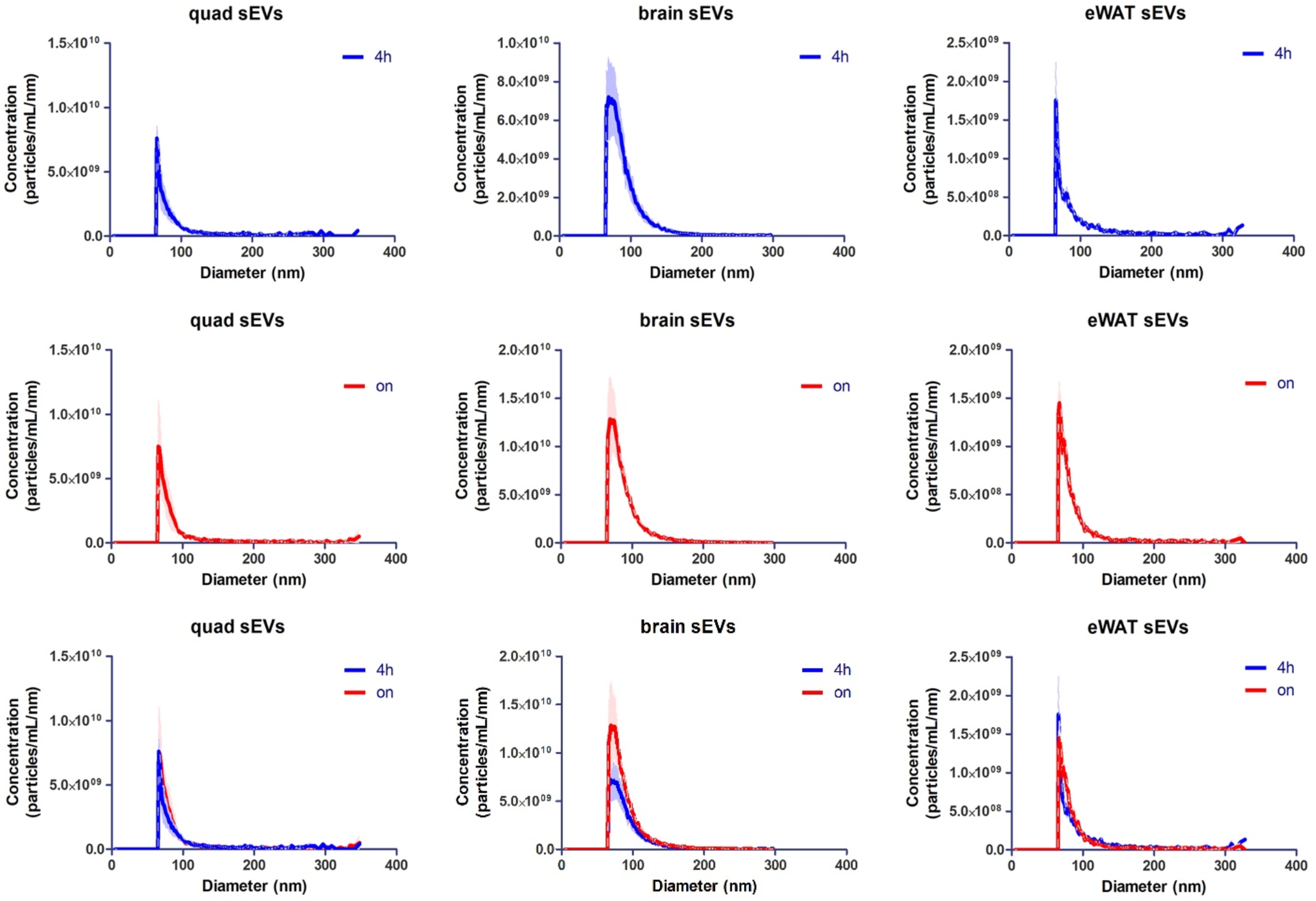
Particle size and concentration of s-EVs-derived from eWAT, Brain and Quadriceps mouse tissues.

**Figure 4:**
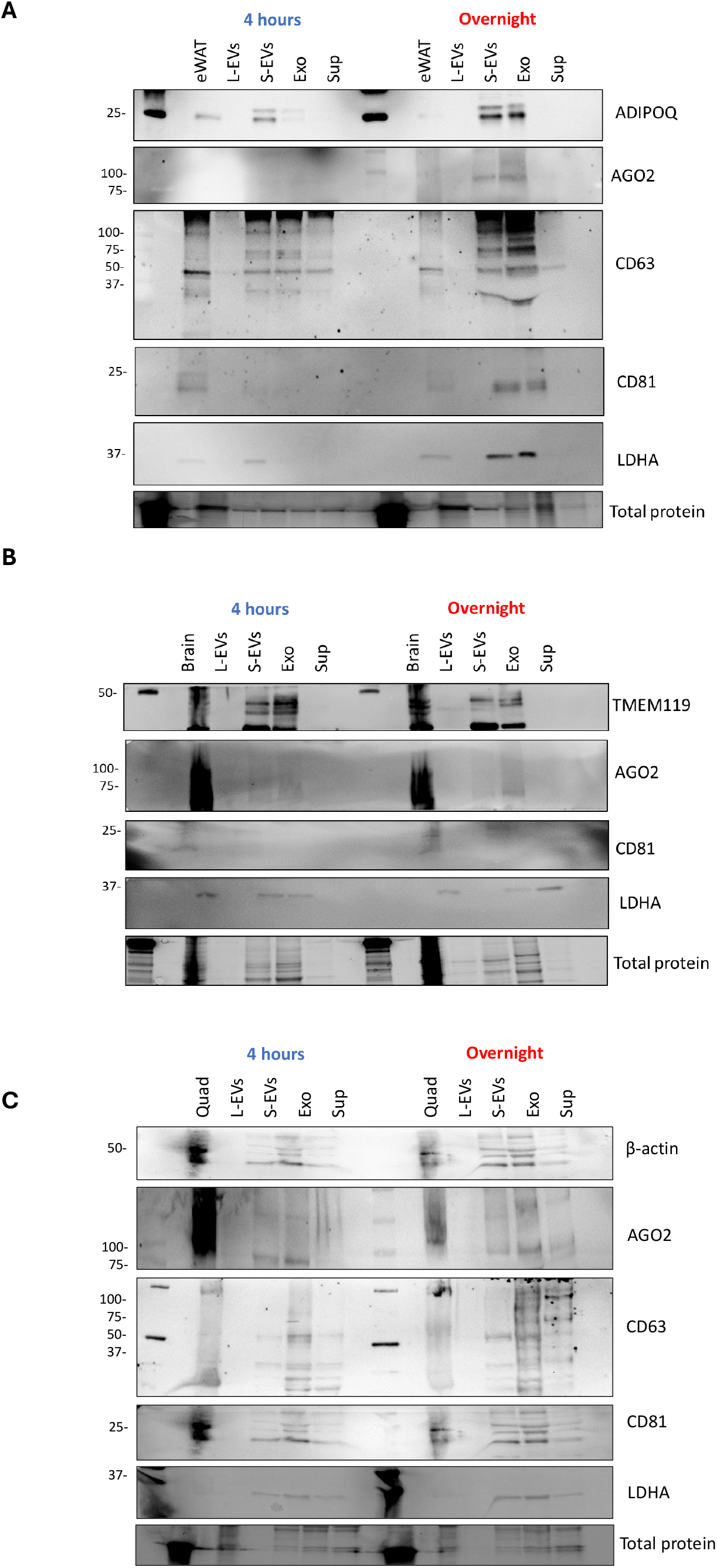
Immunoblot analysis of EV and NVEP markers in (**A**) eWAT, (**B**) Brain or (**C**) Quadriceps mouse tissues.

